# Ecological ubiquity and phylogeny drive nestedness in phages–bacteria networks and shape the bacterial defensome

**DOI:** 10.1101/2025.08.05.668612

**Authors:** Chloé Feltin, Sylvain Piry, Benoit Moury, Lola Chateau, Karine Berthier, Jonathan M. Jacobs, Jules Butchacas, Linda Fenske, Lillian Ebeling-Koning, Theo H. M. Smits, Cindy E. Morris, Clara Torres-Barceló

## Abstract

Identifying the ecological and evolutionary factors that shape phage–bacterial interactions is key to understanding their dynamics in microbial communities. Yet, such interactions remain poorly characterised in plant agroecosystems. Here, we investigate the ecological determinants of the interaction between a highly diverse set of 23 phages isolated from diseased apricot trees and 44 bacterial strains from the *Pseudomonas syringae* species complex collected either from diseased apricot trees, healthy plants or non-agricultural environment. Based on their ecological origin, we expected phages to preferentially infect bacterial strains from the same ecological context, forming modular host-range patterns. Contrary to these expectations, we discovered a significantly nested structure, suggesting generalised infection dynamics rather than local adaptation, primarily driven by the broad ecological dynamics of this pathosystem. Analysis of the bacterial genomes showed that both the profiles of anti-phage defence systems and the distribution profiles of prophages are strongly shaped by bacterial phylogeny. Furthermore, while the number of defence systems showed limited correlation with the breadth of bacterial sensitivity to phages, prophage abundance exhibited a strong, non-linear link with phage virulence. Together, these findings provide an ecological and evolutionary perspective on phage–bacterium infection networks and new insights into a better understanding of the role of phages in agricultural ecosystems.

**Author Summary:** Viruses that infect bacteria, known as phages, are part of microbial communities and influence the abundance, diversity, and traits of their hosts. In an agriculture-related context, they are commonly considered as potential biocontrol agents, but studying the bases of fundamental phage–bacterial interactions may help us better understand the plant microbiome and its applications. Many factors influence these interactions, and identifying which ones matter most remains a challenge. In our study, we investigated how phages from diseased plants interact with bacteria collected from diseased and healthy plants, as well as from surrounding environments. We expected phages to mainly infect bacteria from similar environments, but instead observed that they often infected bacteria regardless of their source. This suggests that phage activity in this system has few barriers, reflecting the wide ecological distribution of their bacterial hosts. We further investigated how bacteria defend against phages by identifying both defence systems and prophages within their genomes and using this information to explore their contribution to bacterial resistance or sensitivity to phages. Together, our findings offer new insights into how phage–bacterium relationships evolve and function in plant ecosystems.

## Introduction

Phages profoundly influence bacterial diversity and population dynamics (1). The impact of phages on bacterial ecology and evolution has been extensively examined in ecosystems like oceans and the human gut (2–7), research in agriculture (8). A deeper understanding of the ecological roles, population dynamics, and interactions of phages with bacterial communities in plant-associated systems, could provide fundamental insights to guide the development of phage-based biocontrol strategies and the sustainable management of microbial communities in agriculture.

Phage interactions with bacteria depend on a multitude of factors. The phage must recognise and infect a host, integrate or replicate, while avoiding or blocking bacterial defence systems (9,10). A range of approaches have been employed to unravel the determinants of phage-bacterial interactions, including ecological and evolutionary analyses of natural populations and experimental evolution (11,12). Mechanistic studies have identified bacterial receptors (13,14), with more recent work focusing on defence systems associated with susceptibility to phage (14–17). For example, *Vibrio lentus* strains with identical phage receptors display varying levels of susceptibility depending on their phage defence elements (18). However, research on prophages and defence systems in phytopathogenic bacteria remains limited and have yet to connect these elements to host range data (19–25).

The complexities of phage-bacterial interactions can be visualised as ecological networks, which exhibit distinct structural patterns, commonly modularity or nestedness (26). Nestedness, a structure in which phages exhibit a gradient of host range breadth and bacteria varying levels of susceptibility, is commonly observed with limited genetic diversity of phages or bacteria (26–28). In contrast, modularity, where phages and bacteria interact within modules but rarely across them, tends to emerge at broader evolutionary or ecological scales, such as when interactions involve hosts, or are examined over extended time periods (11,29). Nonetheless, modules can also evolve rapidly and emerge within species (*e.g*. (12)), although such events may be rare. A recent study introduced statistical methods that incorporate quantitative host–parasite interaction data into ecological networks (30). By considering interaction strength rather than simple infection presence or absence, these approaches provide a more nuanced view of infection patterns, host susceptibility, and phage infectivity, capturing dynamics overlooked by binary analyses.

The *Pseudomonas syringae* species complex is particularly interesting for studying phage-bacterial interactions in agriculture, due to its high genetic diversity and broad ecological distribution across cultivated plants, wild environments and aquatic habitats (31–33). *P. syringae* strains responsible for bacterial canker of apricot trees (*Prunus armeniaca*) are particularly diverse, spanning multiple phylogroups (PG01, PG02, PG03, PG07) (34). It is hypothesized that the pathogenic success of these apricot strains may be driven primarily by general ecological fitness rather than specific virulence factors. These lineages are not apricot-specific but thrive across varied environments, raising questions about the ecological contexts of their phage interactions, whether within the host or external reservoirs (34). Exploring phage infectivity patterns on these strains can reveal how phylogenetic and ecological diversity shapes phage-bacterium dynamics.

Historically, phage host range profiling was used to differentiate *P. syringae* strains (35–39). More recent studies have emphasised the isolation and characterisation of novel *P. syringae* phages and their potential as biocontrol agents (40,41). Most phage host range assays were confined to the isolation host and the intended target, reflecting an applied focus. Others included a broader set of strains, though typically limited to agricultural isolates. For example, phages infecting *P. syringae* pv. *porri* exhibited strict specificity to this variant, having been tested mainly against *porri* strains and a few non-*porri* isolates, and *P. syringae* pv. *actinidiae* phages were largely pathovar-specific, with some extending activity to closely related lineages (42).

We focused on the causal agents of bacterial canker of apricot trees to investigate the structure and ecological drivers of phage–bacterium infection networks. We hypothesised that phages would be locally adapted to their bacterial hosts, leading to a modular infection network structure that could be explained by the presence of prophages and diverse bacterial defence systems. To test this, we addressed the following questions: i) Are phages isolated from diseased apricot trees specifically adapted to resident *P. syringae* compared to closely-related strains from other environmental reservoirs? ii) Do closely-related bacteria isolated from different environments differ in their adaptation to phages? iii) What bacterial genomic traits underlie the observed infection network structure? We conducted an integrative approach utilizing a quantitative host-range assay involving 23 genetically-diverse *P. syringae* phages and 44 *P. syringae* strains. This was coupled with whole-genome sequencing of all bacterial strains for detailed annotation of prophages and defence systems. This comprehensive methodology provides mechanistic insights into how ecological origin and bacterial genomic content shape phage–bacterial interactions.

## Results

### Host range characterization of a diverse panel of phages on P. syringae bacteria

We characterised the phage–bacterial infection network by testing 23 phages against 44 strains from the *P. syringae* species complex (hereafter referred to as *P. syringae* strains) in liquid culture over 48 hours of bacterial growth monitoring. Unlike traditional plaque assays, which primarily assess phage replication on solid media, this quantitative approach measures the extent of bacterial growth suppression in liquid conditions, thereby capturing a complementary aspect of phage–bacterium interactions (9,43). Each interaction was quantified using two complementary metrics: 1/ a continuous variable, corresponding to the percentage of bacterial growth inhibition by phages (0–100%) (S1A Table), and 2/ a binary variable, indicating whether the inhibition was statistically significant (1) or not (0), based on triplicate measurements (S1B Table). Because these values can be interpreted from either the phage or bacterial perspective, we provide clear terminology to distinguish them. From the bacterial perspective, these values represent susceptibility (percentage inhibition) and sensitivity (binary significance of inhibition). Conversely, from the phage perspective, these values reflect virulence (percentage inhibition) and infectivity (binary success of infection).

Overall, the tested bacterial strains showed continuous distributions of susceptibility and sensitivity values, reflecting the diversity of interactions of the panel with phages. On average, bacterial susceptibility to phages was moderate, with a 24% reduction in growth, and strains were sensitive to 12.3 out of the 23 tested phages (53.5%) (S1A Fig), indicating a broader sensitivity than suggested by inhibition levels alone. The most susceptible bacterial strains were DG1a.7, GAW0089, and DG12.6 (all from PG02), with average inhibition levels of 46%, 50%, and 42%, respectively (S1A Table). The most broadly sensitive strains, DG1a.7, 7C, and CCE0067 (all from PG02), were significantly inhibited by 20, 19, and 19 phages, respectively (87%, 83%, and 83% of tested phages respectively) (S1B Table). In contrast, bacterial strains CC1583, DG8.15, and DGA9CS5-8 (from PG10, PG07 and PG10 respectively) showed the highest resistance, with average inhibition percentages of only 3%, 6%, and 6%, respectively (S1A Table). DG8.15 (from PG07) resisted to 17 phages (74% of tested phages) and seven other strains (19B, 41D, CST0086, DGA9CS5-15, FMU-107, and USA0088 from PG01, PG02, PG02, PG10, PG07 and PG02) were resistant to 16 phages (70%) (S1B Table).

From the phage perspective, average bacterial growth inhibition across the strain panel was 23%, reflecting moderate virulence. However, each phage was able to infect an average of 22.4 out of 42 tested *P. syringae* strains (53.3%) (S1B Fig), indicating a relatively broad infectivity despite moderate inhibition levels. The most virulent phages were Ghual01 (65% inhibition), Arace01 (57%), and Draal03 (44%) (S1A Table). The most infecting phages were Nican01, Arace01, and Ghual01 infecting 39 (93% of tested bacteria), 37 (88%), and 34 (81%) strains, respectively (S1B Table). Moreover, Arace01 and Ghual01 were the only phages capable of infecting strains from other species in the *Pseudomonas* genus. Arace01 infected a *P. rhizosphaerae* strain at a low efficiency of plating (EOP) of 0.005, whereas Ghual01 infected a *P. graminis* strain at a moderate efficiency (EOP = 0.017). In contrast, Orimi01, Cygsa01, and Pavpe01 displayed the lowest virulence, with an average bacterial growth inhibition of 0%, 3%, and 4% respectively (S1A Table), while Orimi01, Cygsa01, and Touem01 had the lowest infectivity, infecting only 2 (5% of tested bacteria), 5 (12%), and 11 (26%) strains, respectively (S1B Table).

The relationship between the breadth of infection and the strength of inhibition can reveals underlying evolutionary dynamics, including trade-offs in both phages and bacteria (44,45). When considering only significant inhibition interactions, we detected a positive correlation between the phage virulence and the phage infectivity (Spearman rho = 0.56; p = 0.01; S9C Fig), indicating that phages with broader host ranges tend to exhibit higher average virulence not supporting the presence of a trade-off. However, this interpretation should be viewed with caution, as some degree of metric dependency may persist. In contrast, no correlation was observed between bacterial susceptibility and bacterial sensitivity (Spearman rho = 0.24; p = 0.12; S9D Fig), reflecting greater variability among bacterial strains in their response to phage infections and suggesting the potential for trade-offs in bacterial defensive strategies.

These findings reveal substantial variability in phage virulence and infectivity, as well as bacterial susceptibility and sensitivity, with over than half (53%) of combinations resulting in significant growth inhibition. While some phages exhibit a broad host range, extending even beyond *P. syringae*, others are highly specialised, only infecting a few strains, reflecting a complex interplay between phage infectivity and bacterial resistance.

### Nested structure of the phage-bacterial interaction network and association with bacterial traits

The structure of the phage-bacterial infection network was analysed using multiple algorithms to assess its nestedness and modularity. Given the diversity of both phages and bacterial hosts, spanning 13 phage genera and four phylogroups of *P. syringae*, and the ecological origins of phages (diseased apricot trees) and bacteria (diseased apricot trees, healthy plants and other environmental reservoirs), we hypothesised a modular structure type. However, the network representing bacterial growth inhibition percentage by phages displayed a strong nested structure, as indicated by scores of 0.48 and 0.44, on a scale varying from 0 to 1, with the *WINE* and *wNODF* algorithms, respectively (Table 1). To determine the statistical significance of the nested pattern, the network was compared to null models, which are simulations differing in the constraints applied during randomization, ranging from unconstrained (S, N) to fully constrained (B). Nestedness was highly significant (p < 0.01) against nearly all null models, except null model B, which was shown to be excessively conservative (30).

**Table 1.** Analysis of nestedness and modularity of the bacterial growth inhibition network combining 44 strains of the *P. syringae* species complex and 23 phages.

| Analysis | Algorithm | Score<br>(max.<br>= 1.0) | Null model |  |  |  |  |  |  |
| --- | --- | --- | --- | --- | --- | --- | --- | --- | --- |
|  |  |  | B | N | C1 | R1 | S | C2 | R2 |
| Nestedness | wNODF | 0.44 | 0.44 | 0 | 0 | 0 | 0 | 0 | 0 |
|  | WINE | 0.48 | 1 | 0 | 0.01 | 0 | 0 | 0 | 0 |
| Modularity | spinglass | 0.01 | 1 | 1 | 1 | 1 | 0.07 | 0.29 | 0.18 |
|  | edge<br>betweenness | 0.09 | 0 | 0.02 | 0 | 0 | 1 | 1 | 1 |
|  | fast greedy | 0.24 | 0 | 0 | 0 | 0 | 1 | 0.81 | 0.02 |
|  | leading<br>eigenvector | 0.14 | 0 | 0 | 0 | 0 | 1 | 1 | 0.94 |
|  | beckett | 0.25 | 0 | 0 | 0 | 0 | 1 | 0.01 | 1 |

Null models are described in Moury *et al.* (30). For each null model and each algorithm, the frequency of simulations, among 100, showing a strictly higher nestedness (or modularity) score than that of the actual matrix is indicated. Significantly nested or modular networks ( $p \leq 0.05$ ) are indicated in red.

Modularity analyses differed depending on the algorithm (Table 1). With the *spinglass* algorithm, selected for its good performance in terms of false positive rate (30), the modularity score was low (0.01, on a scale varying from -0.5 to 1) and was not significant, whatever the null model. The four alternative modularity algorithms produced slightly higher scores (0.09 < modularity < 0.25) and showed significant modularity (p < 0.01) with both null models C1 and R1. These four algorithms show lower performances than *spinglass* in terms of false positive rates, but combined with null models C1 and R1, they show generally acceptable performances in terms of false positive and false negative rates (30).

These results suggest globally a slight degree of modular organisation within the network. The strong and significant nestedness and the weak modularity were further supported by the analysis of the binary data representing the significance of bacterial growth inhibition (S2 Table).

To assess whether the nested structure of phage–bacterial networks is shaped by bacterial evolutionary history, we first tested for a phylogenetic signal in bacterial sensitivity summed by bacteria (Fig 1). The analysis revealed a strong phylogenetic signal (λ = 1.0; LR (λ = 0) = 48.26; p < 0.01), indicating that closely-related bacterial strains tend to be infected by a similar number of phages. To explore this pattern further, we evaluated the effect of phylogroup identity using both a Generalised Linear Model (GLM) with the bacterial sensitivity summed by bacteria (Fig 1C) and a PErmutational Multivariate ANalysis Of VAriance (PERMANOVA) on bacterial susceptibility profiles (S2A Fig). Both analyses confirmed a significant influence of phylogroup on bacterial susceptibility, explaining almost 10% of the variance (GLM: p < 0.01; PERMANOVA: R² = 0.09; p < 0.01). In contrast, ecological origin had no significant effect in either test (GLM: p = 0.87; PERMANOVA: R² = 0.05; p = 0.29). Specifically, PG02 strains emerged as the most sensitive to phages, being infected on average by 14.6 phages and showing the highest susceptibility, with 20.7% growth inhibition. PG01 strains showed intermediate sensitivity being infected by 10.4 phages and an average of 14.6% inhibition, followed closely by PG10 strains with 15.3% inhibition. In contrast, PG07 strains were the most resistant overall, infected on average by only 8.4 phages and displaying the lowest susceptibility with 8.73% inhibition.

**Fig 1.**
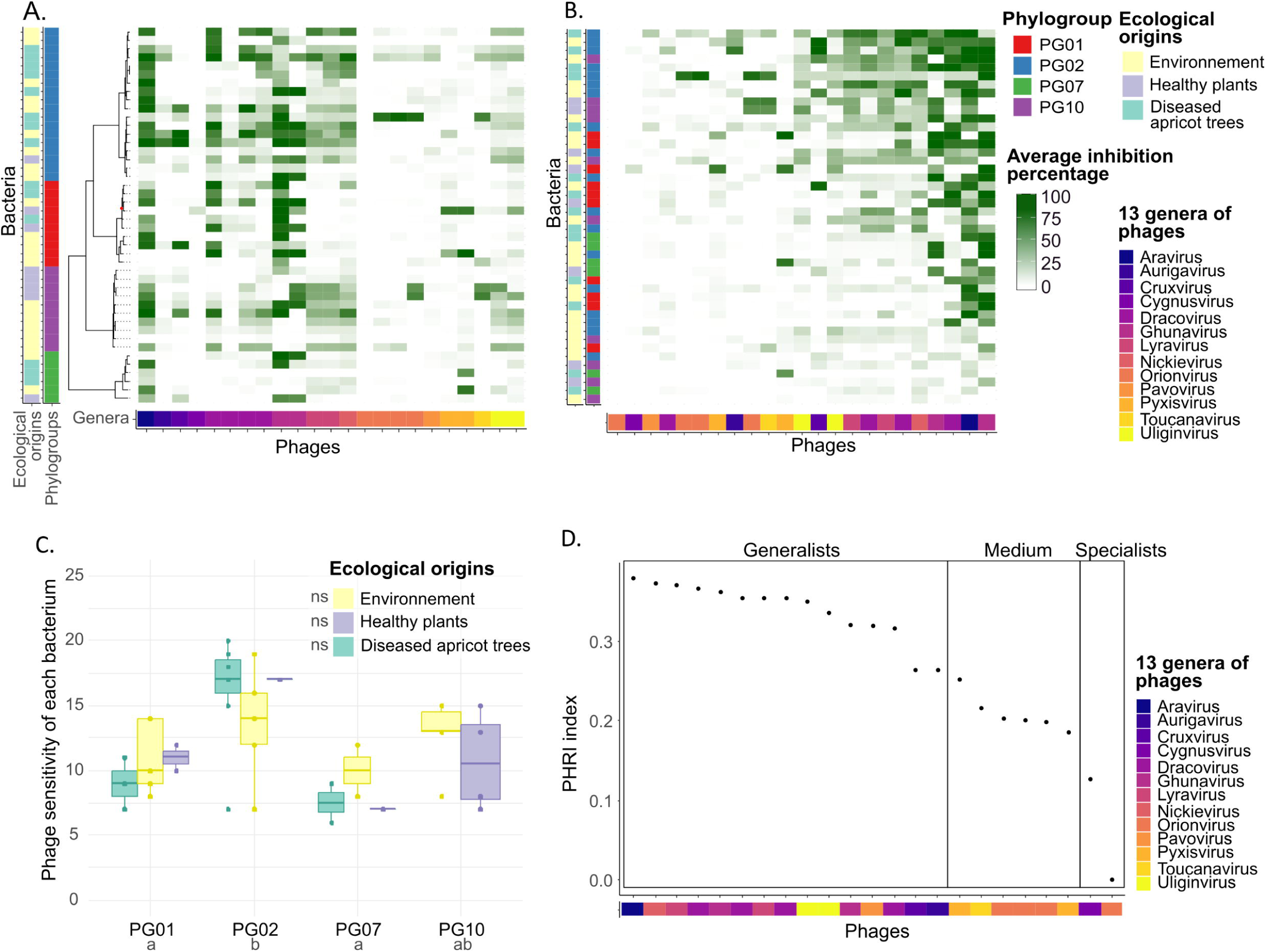
Networks of percentages of growth inhibition of 44 P. syringae species complex strains (rows) by 23 phages (columns). (A) Taxonomic organisation of the network presented on a maximum likelihood phylogenetic tree based on bacterial core-genomes and phage genera. A node with a maximum likelihood support value below 0.90 is marked with a red dot. (B) The same network after row and column permutations showing its nested pattern, with phylogroups and ecological origins of *P. syringae* strains displayed on the right side of panels A and B. (C) Bacterial sensitivity to phages by bacterial phylogroup and ecological origins. Letters indicate Tukey post-hoc test results from a generalised linear model (GLM) with a Poisson distribution; ns indicates non-significant differences (p > 0.05). (D) Phylogenetic Host Range Index (PHRI) across phage genera.

The broad taxonomic diversity of the phages studied here, covering 13 genera, includes both specialists and generalists. To characterise phage host range, we calculated the Phylogenetic Host Range Index (PHRI), which incorporates the number, the evenness, and the phylogenetic diversity of host bacteria (46). The PHRI values ranged from 0 (Orimi01, a specialist phage) to 0.38 (Arace01, a generalist phage), forming a gradient that reflects the nested structure of the phage-bacteria interaction network (Fig 1D). Based on PHRI classification, 9% of phages were categorised as specialists, 65% as generalists, and 26% as having an intermediate host range (Fig 1D). At the genetic level, significant differences in virulence were observed between phage genera (GLMM; p < 0.01). The generalist genus *Aravirus* exhibited the highest virulence (52.8%), while the specialist genera *Cygnusvirus* and *Orionvirus* displayed markedly lower virulence levels, averaging 1.1% and 4.2%, respectively (S2B Fig). Additionally, phages with similar host infection profiles clustered by taxonomy, with those sharing >70% nucleotide identity (same genus) explaining 63.5% of the variation in virulence (PERMANOVA: R² = 0.64, p < 0.01), indicating genus-level host specificity (Fig 1A). For example, the two phages from the *Lyravirus sulafat* species and the *Uliginvirus* genus exhibited highly similar infection host profiles.

These results demonstrate that the phage–bacterial interaction networks are predominantly nested within the investigated panel of strains, with minimal modularity suggesting broad host range overlap among phages rather than local adaptation based on ecological origin. Bacterial susceptibility is phylogenetically constrained, with PG02 strains being highly sensitive and PG07 strains resistant. Additionally, although more than half of the phages were generalists, genus-level similarity drives host-range patterns.

### Defence systems: profile and association with ecology and phylogeny

The 44 strains of *P. syringae* analysed in this study were examined for their accessory genome content, with a particular focus on their repertoire of defence systems to counteract mobile genetic elements such as phages. Across these strains, a total of 48 defence system types and 86 subtypes were identified (Fig 2A). The most abundant defence subtypes included Septu (*n* = 81), type IV Restriction-Modification (RM) (*n* = 37), and Gabija (*n* = 31) (Fig 2A). In contrast, 31 subtypes were sporadically detected across the bacterial genomes, including rare defence systems such as three Abi subtypes (AbiH, AbiJ, and AbiO), two Asv subtypes (AsvI and AsvII), and three DTR subtypes (DTR6, DTR8, and DTR3) (Fig 2A). Defence systems are present at an average rate of 9.93 per strain, with two strains (AF0015 and P3.01.09C09, both from PG01) harbouring the highest number (15 systems), and four strains (DG1A_7 from PG02, CSZ0293, SZ0119, and SZB0008 from PG10) the lowest number (5 systems) (Fig 2A). Each defence system can be classified into different functional categories to better characterise its role. Specifically, 38.4% of defence system subtypes are associated with abortive infection, 24.4% with nucleic acid degradation, and 37.2% remain unclassified (Fig 2A).

**Fig 2.**
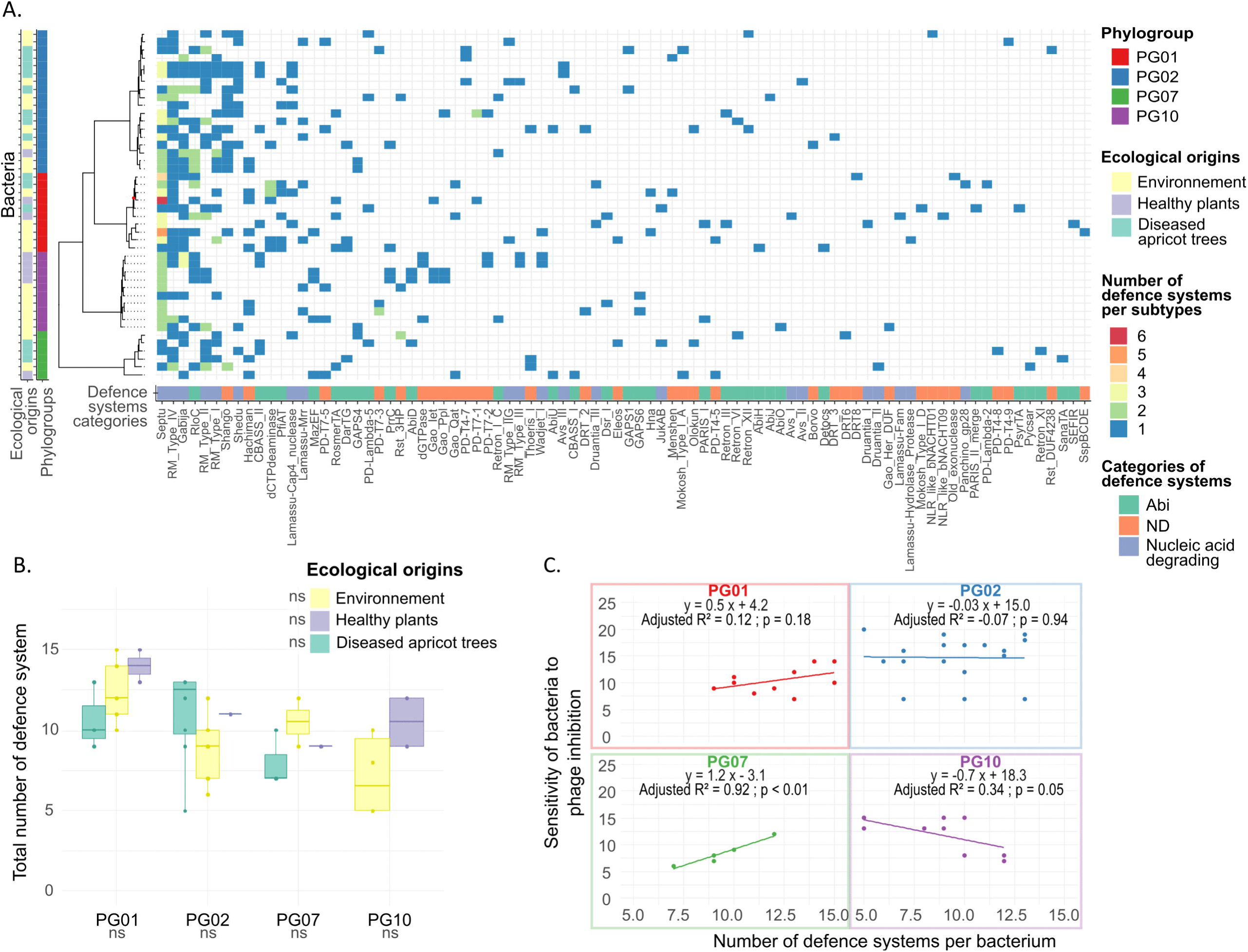
Defence system distribution in bacterial genomes and relationship with bacterial sensitivity to phages. (A) Matrix showing the number of defence systems present in the 44 *P. syringae* strains, organised on a maximum likelihood phylogenetic tree based on their core-genomes. A node with a maximum likelihood support value below 0.90 is marked with a red dot. The phylogroup affiliation of each bacterial strain is indicated on the left using colours, and defence systems are represented by subtypes with colours on the right indicating their respective functional category. (B) Number of defence systems per bacterial strain according to their phylogroups and ecological origins. ns: not significant (p > 0.05), according to post-hoc Tukey tests from a Generalised Linear Model with a Poisson distribution. (C) Bacterial sensitivity to phages as a function of the number of defence systems, with linear regressions performed within each phylogroup.

To assess whether the distribution of defence systems across bacterial genomes was influenced by phylogeny, we tested for a phylogenetic signal in the total number of encoded systems per genome. A strong signal indicated that closely related strains tend to harbour similar numbers of defence systems (λ = 1.0; LR test (λ = 0): 44.18; p < 0.01) (S3 Fig), as described in a previous study (24). Phylogroup significantly explained variation in defence system profiles (PERMANOVA, R² = 0.14, p < 0.01), with a weaker effect from ecological origin (R² = 0.06, p < 0.01), while growth rate of the bacteria had no significant impact (R² = 0.02, p = 0.51) (Fig 2A). Post-hoc comparisons based on this analysis revealed that strains from PG10 displayed significantly distinct defence system profiles compared to those from PG02 and PG07, while strains isolated from healthy plants exhibited distinct defence system profiles compared to other ecological origins. However, none of these factors significantly affected the total number of defence systems, although the effect of phylogroup approached significance (GLM: p = 0.06) (Fig 2B). This phylogroup trend is supported by a larger dataset of 590 *P. syringae* genome assemblies, where PG01 strains encode the highest number of defence systems, while PG07 and PG10 harbour some of the lowest counts (24).

While closely-related strains tend to possess similar numbers of defence systems, we further investigated whether they also share the same subtypes of defence systems. To this end, we calculated the percentage of shared defence system subtypes between each pair of bacterial strains, where 100% indicates complete identity in the sets of subtypes. A weak but significant positive correlation was observed between the percentage of shared defence system subtypes and the genome-wide Average Nucleotide Identity (ANI) among bacterial strains (GLMM; estimate = 0.015; p < 0.01) (S4 Fig). This relationship suggests that some defence systems are partly inherited through vertical transmission, reflecting the influence of phylogeny on their distribution. Furthermore, only 1.14% of the identified defence systems were linked to prophages within bacterial genomes, indicating that the vast majority are located outside prophage regions. This low prevalence indicates that while defence systems may spread horizontally among bacterial populations, their dissemination is likely mediated by mobile genetic elements other than prophages.

Defence systems may impose varying energetic costs on bacterial cells, potentially influencing their allocation of resources (47,48). In strains of *P. syringae*, defence systems related to nucleic acid degradation comprised, on average, 54% of the total defence systems, while abortive infection systems, which are considered more metabolically costly (49,50), constituted an average of 25% (S3 Fig). We then revealed that the proportion of defence system categories were significantly shaped by phylogroup, explaining 16% of the variance (PERMANOVA; R² = 0.16; p = 0.03), and growth rate (R² = 0.07; p = 0.04), but not ecological origin (R² = 0.08; p = 0.11). Pairwise comparisons based on this analysis showed that strains from PG07 had significantly different defence category profiles compared to those from PG01 and PG02.

Our results revealed that the defence arsenal profiles in *P. syringae* are shaped by phylogeny and slightly by ecological origin, consistent with partial vertical inheritance of defence system profiles. Overall, these findings underscore the complexity of bacterial defence strategies and indicate that multiple evolutionary and ecological factors influence phage resistance.

### Prophages: profile and association with ecology and phylogeny

We identified 88 prophages among the 44 genomes of *P. syringae*. On average, each strain carried two prophages, with one strain (FMU-107) harbouring five prophages and 18 strains (41%) containing a single cryptic prophage (Fig 3A). Among these prophages, 36% were classified as high quality (complete or nearly complete according to CheckV software), 39% as medium quality, and 25% as low quality. Prophage sizes ranged from 12.0 kb to 89.8 kb, with an average size of 32.2 kb. The size distribution exhibited three peaks at approximately 18 kb, 44 kb, and 57 kb. To classify these prophages taxonomically, we used VIRIDIC, which identified 65 novel phage species and 42 novel genera (S3 Table), while proteomic tree analysis with Viptree revealed six distinct prophage clusters, corresponding approximately to family-level taxonomic divisions (Fig 3A) (51). One of these, the CL4 family-like prophage, was found in all 44 *P. syringae* strains, which span four distinct phylogroups (Fig 3A). CL4 family-like prophage is a tailocin, specifically an R-type syringacin, a bacteriocin derived from a bacteriophage (52,53). These tailocins are widely distributed across the *P. syringae* species complex and are thought to play a key role in inter-bacterial competition by enabling strains to kill closely-related competitors (54,55). Moreover, we found that the taxonomy of the CL4 family-like prophage mirrors the phylogeny of their bacterial hosts across 68 strains from 10 phylogroups (including the 44 strains analysed in this study), with a significant cophylogenetic congruence signal (Procrustes Approach to Cophylogeny (PACo): residual sum of squares = 0.15; *p* < 0.01) (S5 Fig).

**Fig 3.**
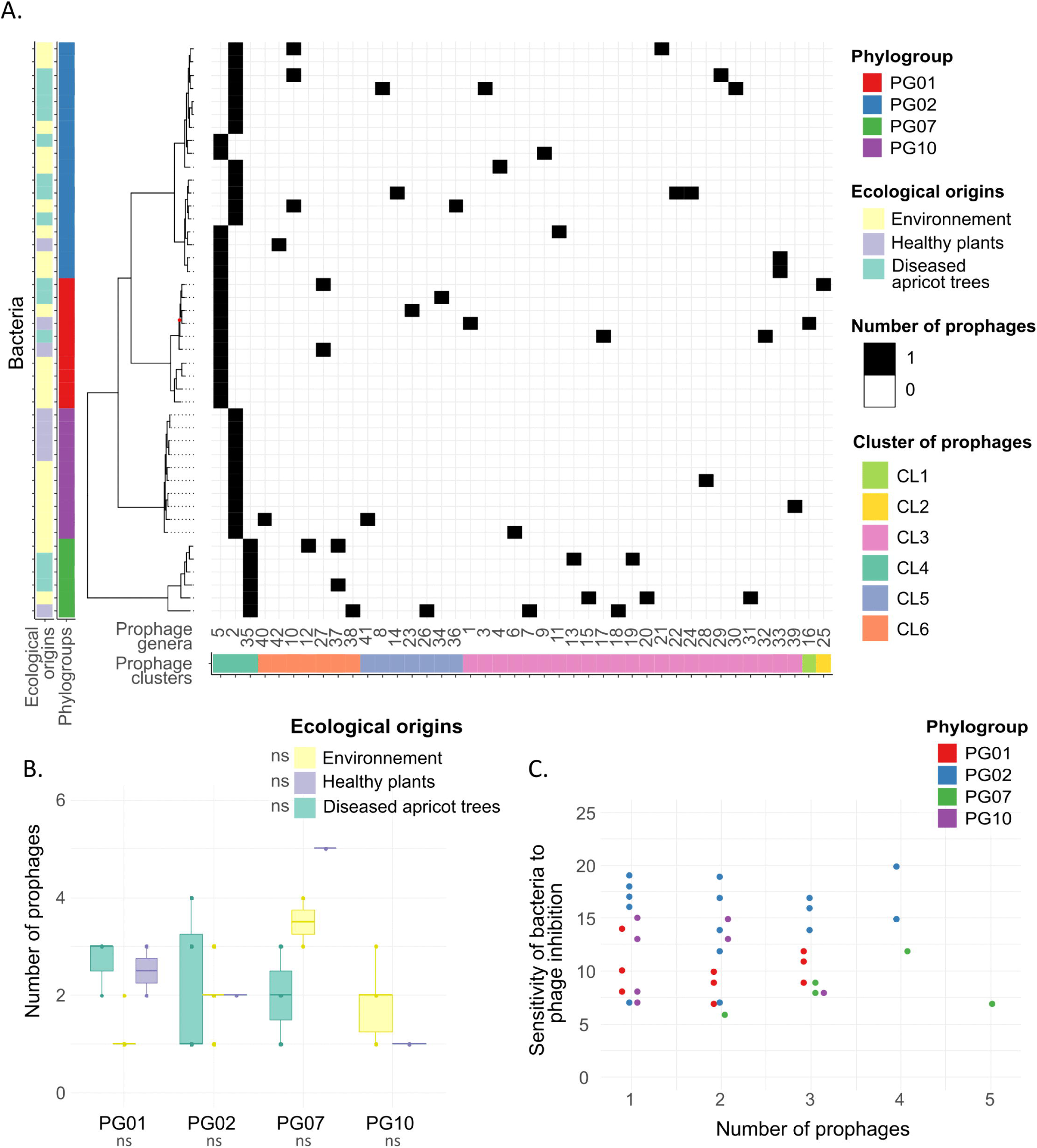
Diversity and distribution of prophages in bacterial genomes and relationship with bacterial sensitivity to phages. (A) Distribution of prophages in the genomes of the 44 *P. syringae* strains, categorised by genus and family-like cluster (VipTree). Bacterial strains are organised based on a maximum likelihood phylogenetic tree based on their core-genomes, with their phylogroups and ecological origins indicated on the right. A node with a maximum likelihood support value below 0.90 is marked with a red dot. (B) Abundance of prophages per bacterial strain according to their phylogroups and ecological origins. ns: not significant (p > 0.05), according to post-hoc Tukey tests from a Generalised Linear Model with a Poisson distribution. (C) Bacterial sensitivity to phages as a function of the number of prophages, coloured by phylogroups.

The abundance of prophages (*i.e.* the number of prophages per bacterial genome) and the prophage distribution profiles in bacterial genomes are strongly shaped by phylogeny. A significant phylogenetic signal was detected in prophage abundance (λ = 1.0; LR (λ = 0) = 46.48; p < 0.01), indicating that closely related strains tend to harbour similar numbers of prophages (Fig 3B). Phylogroup significantly explained 16% of variation in prophage distribution profiles (PERMANOVA, R² = 0.16; p < 0.01), with a weaker effect from ecological origin (R² = 0.07; p < 0.01), while growth rate had no significant impact (R² = 0.03; p = 0.09) (Fig 3A). All distinct phylogroups, as well as ecological origins, exhibited significantly different prophage distribution profiles, supporting the idea that lineage and ecological context influence the genomic landscape of prophages. However, none of these factors significantly affected the abundance of prophages (GLM: phylogroup, p = 0.27; ecological origin, p = 0.92; growth rate, p = 0.30), despite some observed variation. Indeed, PG07 strains showed the highest prophage abundance, nearly double that of PG10 strains (Fig 3B). However, this pattern contrasts with findings from broader dataset (590 *P. syringae* genome assemblies), in which PG02 strains encoded among the highest numbers of prophages, PG07 among the lowest, and PG01 and PG10 showed intermediate levels (24). In contrast, differences across ecological origins were minimal (Fig 3B).

In conclusion, the comprehensive analysis of prophages within the *P. syringae* species complex revealed a diverse array of complete and incomplete prophages integrated into the genomes. The prophage distribution profiles are strongly explained by bacterial phylogeny and ecological origins of the host strains.

### Contribution of bacterial defence systems and prophages to bacterial sensitivity to phages

Since bacterial defence systems target phages (50), a negative correlation is expected between their number and bacterial sensitivity to phages. To test this, we examined the correlation between bacterial sensitivity and the number of encoded defence systems, while accounting for phylogenetic background. When considering all bacterial strains together, no overall correlation was identified (p = 0.36) with a phylogroup effect (p < 0.01) and no effect of ecological origins (p = 0.88). For both PG01 and PG02, no significant correlation was detected (linear regression for PG01: adjusted R² = 0.12; p = 0.18; for PG02: adjusted R² = – 0.07; p = 0.94) (Fig 2C). In contrast, a negative correlation was detected in PG10, supporting our expectation (linear regression for PG10: adjusted R² = 0.34; p = 0.05) (Fig 2C). In contrast, PG07 exhibited an unexpected positive correlation, suggesting that strains with a higher number of defence systems were more sensitive to phages (linear regression for PG07: adjusted R² = 0.92; p < 0.01) (Fig 2C).

Although prophages can contribute to bacterial immunity against phage infection (56), we found no significant association between prophage abundance and bacterial sensitivity to phages (p = 0.82), and neither phylogroup (p = 0.31) nor ecological origin (p = 0.80) had a detectable effect (Fig. 3C). Moreover, no correlation was observed between prophage abundance and the number of defence systems encoded in bacterial genomes, suggesting that these systems do not provide protection against prophage integration or maintenance (linear regression; adjusted R² = –0.02; p = 0.64).

### Influence of multiple bacterial and phage traits on bacterial growth inhibition

Having identified several bacterial and phage factors that individually influenced the bacterial growth inhibition, we next sought to integrate these explanatory variables into a unified framework to gain a more comprehensive understanding of phage–bacteria interaction dynamics. The primary model included prophage abundance and the total number of defence systems per strain as bacterial traits associated with phage resistance, alongside phage- and bacterium-related traits examined in this study. We also tested an alternative model incorporating PCA-derived variables related to bacterial defensome, but it did not improve model performance (Akaike Information Criterion (AIC) = 16.70) compared to the initial model (AIC = 16.69). Therefore, we retained the first model for its greater interpretability.

Consistent with the above findings, bacterial phylogeny and phage taxonomy emerged as the strongest explanatory variables for inhibition percentage, with an importance weight of 1.0 across the 256 models (Table 2). In addition, a variable indicating whether the phage had been isolated using one of the tested bacterial strains was significantly associated with inhibition level. Specifically, phages isolated from strains within the tested host range exhibited a slightly lower inhibition capacity (mean = 15.1%) compared to those isolated from external strains (mean = 17.7%).

**Table 2.**
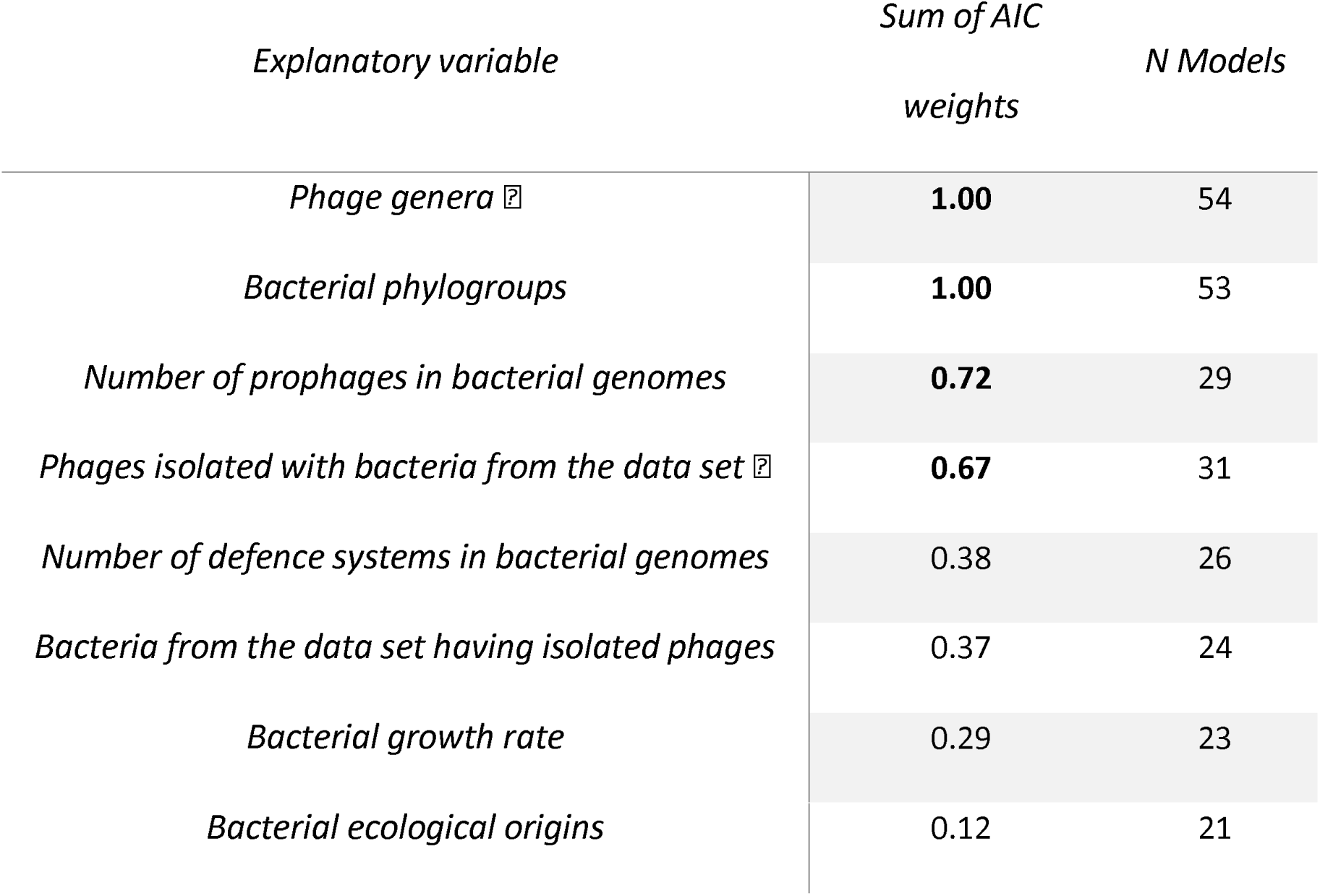
Relative importance of explanatory variables influencing bacterial growth inhibition by phages.

Sum of AIC weights was computed from model averaging across the top-ranked models whose cumulative weights reached ≥ 0.95, selected from a full set of 256 candidate models generated by multi-model inference. The relative importance of each variable corresponds to the sum of AIC weights across all retained models in which it appears. Bold values indicate variables with high relative importance (sum of weights ≥ 0.60), selected as significant predictors in the analysis. N Models: Number of retained models in which each variable appears. Variables related to phages are marked with the symbol V; bacterial variables are unmarked.

The model identified the total number of prophages in bacterial genomes as a significant predictor of inhibition of bacterial growth (sum of weights = 0.72; Table 2). Inhibition levels varied nonlinearly with prophage abundance: inhibition levels were 15.6%, 14.7%, 17.5%, 28.4%, and 10.9% for strains carrying 1 to 5 prophages, respectively. While prophage abundance did not predict bacterial sensitivity, it appeared to influence bacterial susceptibility, suggesting that prophage-rich bacteria may be more susceptible to highly-effective phages. In contrast, the total number of defence systems did not significantly explain bacterial growth inhibition (sum of weights = 0.38; Table 2), despite its correlation with the bacterial sensitivity for some phylogroups.

Overall, these findings highlight that the primary determinants of the bacterial growth inhibition by phage infection within the *P. syringae* species complex are bacterial phylogeny and phage taxonomy rather than bacterial ecological origins. The integration of multiple explanatory variables within a unified modelling framework further revealed that prophage abundance, but not the total number of defence systems, significantly contributes to variations in inhibition levels. This suggests that while prophages may not act as strict barriers to infection, their presence can modulate the efficacy of phage infection. Together, these results emphasize the complex and multifactorial nature of bacterial susceptibility to phages, shaped by both evolutionary history and prophage-mediated genomic landscapes.

## Discussion

Our study uncovered a strong nested structure in the phage–bacterial infection network involving genetically-diverse *P. syringae* strains and phages. This discovery is in strong contrast with the modular structure frequently reported (11,29). This pronounced nestedness likely reflects the ubiquitous nature of the highly diverse *P. syringae* complex, mirrored by a phage community ranging from specialists to generalists in host range. While bacterial phylogeny had a marked influence on the nestedness of the infection network, we found no evidence of local phages-bacteria co-adaptation. Defence systems and prophages contributed to phage resistance, but their distribution was largely shaped by bacterial phylogeny and ecological origin, rather than phage pressure alone. Together, these findings highlight how evolutionary and ecological forces shape phage–bacterial infection networks in natural environments.

### Ecological origin and phylogenetic influences on bacterial sensitivity to phages

The observed nested infection network is unexpected when we consider the high genetic diversity of the phages (13 genera, 17 species) and bacteria (4 phylogroups corresponding to species levels and 7 clades), as well as the varied ecological origins of the bacterial strains included in this study. Such a diversity of organisms could be expected to reveal local adaptation, where phages perform better on sympatric than allopatric bacterial hosts, in line with what has been demonstrated previously (57,11). However, our findings suggest that phage–host interactions in this pathosystem transcend ecological boundaries. Phages isolated from apricot tree soil infected bacterial strains from both diseased orchards and aquatic environments with comparable efficiency, and bacterial origin did not broadly influence bacterial sensitivity/susceptibility to phages. This absence of modularity, combined with a lack of ecological signal in host range, points to broader ecological connectivity and population flow within the *P. syringae* species complex. These results align with previous work (58) showing that *P. syringae* population structures are shaped more by fluctuating ecological dynamics than by geographic isolation, likely due to documented exchanges between agricultural and aquatic environments (59). Overall, the nested structure observed here implies that interaction dynamics in this system are maintained by widespread generalism and ongoing ecological mixing, rather than niche-specific adaptation.

Bacterial phylogeny plays a central role in shaping both the composition of defence systems, the distribution of prophages, and the sensitivity of *P. syringae* strains to phages. In our study, PG02 strains, known for their ubiquitous presence across habitats and for containing strains capable of infecting a broad range of plant species (60,33), were also the most sensitive to phages, carried a moderate number of defence systems, and encoded one of the highest prophage loads (24). In contrast, PG07 strains, which are predominantly associated with diverse environmental reservoirs (60), were the most resistant to phages despite encoding relatively few defence systems and prophages. PG01 strains, mainly isolated from diseased plants (60), showed intermediate bacterial sensitivity to phages, possessed the greatest number of defence systems, and encoded a moderate prophage load (24). Finally, PG10 strains, exclusively environmental, showed moderate bacterial sensitivity to phages, low numbers of defence systems, and a moderate prophage count (24). These findings suggest a potential trade-off between ecological ubiquity and bacterial sensitivity to phages, a hypothesis that requires direct experimental investigation. In a similar vein, a recent study reported that effector gene counts in *P. syringae*, which are key to pathogenicity, positively correlate with phage defences and show partial conservation within phylogroups (24).

Phylogroups show consistent patterns in defence system profiles, with closely-related strains sharing portions of their defence arsenals, while ecological origins accounted for only minor similarities. Strains within the same phylogroup also tend to retain consistent features in their defensome, including the number and functional categories of defence systems. Collectively, these findings suggest that each phylogroup may adopt a common anti-phage strategy, implemented through distinct combinations of defence systems, mainly acquired by horizontal gene transfers (50,61). For strains belonging to phylogroup PG10, we observed a negative correlation between the number of defence systems and sensitivity to phages, suggesting a protective effect from accumulating diverse systems in their genome (17). Conversely, strains in PG07, although generally the most resistant to phages, showed a positive correlation between bacterial sensitivity to phages and defence systems. This counter-intuitive pattern could reflect historical predation by phages, where strains subjected to intense phage attack would have accumulated defence systems (62). Alternatively, the apparent lack of protection may arise because these systems target phages or mobile genetic elements (MGEs) other than those tested here. However, it is important to interpret these trends cautiously, as they are based on relatively small sample sizes (n=6 for PG07; n=10 for PG10), and our understanding remains partial given the continual discovery of novel defence systems (63). Although the sensitivity of specific phylogroups to phages may be correlated to their defence systems, the overall contribution of these systems to phage resistance in *P. syringae* strains appears limited and may depend on the activity of particular defence mechanisms (64).

### Limitations and alternative approaches for understanding phage-bacterium interactions

Liquid culture assays offer the advantage of measuring a proxy for phage virulence by continuously monitoring bacterial growth kinetics in a high-throughput format. This automated approach enhances reproducibility, allows the simultaneous processing of a large number of samples, and enables the use of diverse metrics to quantify phage-induced inhibition of bacterial growth (65), including host-range, and potential indicators of bacterial resistance to phages (9,43). In our dataset, certain phage–bacterium combinations exhibited signs of evolutionary rescue (when a bacterial population initially suppressed by phages recovers through the growth of resistant mutants) (66) (S11 Fig). Future studies could build on these findings by examining their genetic co-evolution through temporal sampling and genomic analyses. Nevertheless, we acknowledge that the phenomenon of lysis from without (referring to bacterial lysis without phage replication, typically resulting from high phage adsorption rates or abortive infections) cannot be entirely ruled out in our dataset (67). Follow-up experiments measuring phage titres, bacterial population dynamics, or adsorption efficiency would be valuable to confirm the underlying mechanisms of bacterial suppression. Such analyses would also refine our understanding of the role of defence systems during phage infection (14,17). In particular, quantifying phage adsorption rates could help discriminate between pre- and post-adsorption resistance: successful adsorption without subsequent lysis would suggest the involvement of intracellular defence mechanisms, whereas failed adsorption would point to surface receptor incompatibility as the main barrier to infection.

Analysis of bacterial membrane receptors recognised by phages is therefore essential in interpreting host range patterns (68,12). Such factors have enabled accurate prediction of phage-bacteria interactions at the strain level in *E. coli* (14). However, receptors for phages infecting *P. syringae* remain poorly characterised compared to other bacterial models, with only three distinct receptors identified in the *P. syringae* phage receptor database compared to over 50 in *E. coli* (69). Increasing research on phage–phytopathogen interactions will help to close this knowledge gap and improve our understanding of the molecular basis underlying the observed interaction patterns.

A complementary strategy to uncover bacterial genetic determinants of sensitivity to phages and resistance phenotypes involves genotype–phenotype association analyses, including approaches based on gene presence–absence matrices, phylogenetic trait correlation methods and genome-wide association studies (GWAS). This latter method has recently been applied to 80 *P. aeruginosa* strains, identifying host genes that influence susceptibility to two phages, particularly those related to flagella and LPS/O-antigens associated with lipopolysaccharide biosynthesis (70). While our dataset of 44 *P. syringae* genomes may provide a sufficient basis for GWAS, challenges remain. The high genomic diversity of the *P. syringae* species complex and the large number of phages tested (n = 23), each potentially interacting with different receptors or resistance pathways, add considerable complexity. Future studies may address this by focusing on specific phage-host combinations or by applying comparative genomics approaches that can help disentangle the genetic basis of resistance across diverse strains.

### Applied perspectives and future directions

From an applied perspective, we identified several generalist phages capable of infecting a wide range of *P. syringae* species complex strains. Understanding the molecular basis of their generalism could provide important phage evolutionary clues, *e.g.,* whether generalism involves flexible receptor-binding proteins, the ability to evade diverse bacterial defence systems, or other mechanisms. Many *P. syringae* strains are non-pathogenic members of the plant microbiome, highlighting the need for targeted phage cocktails that selectively control pathogenic strains while preserving beneficial microbes. Such specificity offers a more sustainable and ecologically-balanced approach to biocontrol compared to using broad-spectrum generalist phages (71). This exemplifies how fundamental ecological understanding of phage–bacterium infection networks can directly inform and enhance the development of effective biocontrol strategies.

To further test our hypothesis that ecological connectivity shapes the nested structure of phage–*P. syringae* interactions, future studies should isolate phages from diverse non-agricultural environmental, such as water and healthy plants to include cases of true sympatry. These newly isolated phages should then be tested against the same bacterial strains used in our study, or against new strains from corresponding habitats. If similar nested network structures emerge, it would strengthen the idea that phage and *P. syringae* populations continuously migrate between agricultural and non-agricultural environments, forming an ecologically-connected meta-community. Moving forward, several key research priorities should guide future investigations: verification of the role of specific defence systems through genetic manipulation like gene deletion and heterologous expression; systematic investigation of receptor usage by our studied phages to reveal molecular determinants of host range; and strategic expansion of the bacterial strain collection to improve phylogenetic and ecological coverage, with particular focus on understudied phylogroups. These complementary approaches would collectively strengthen our understanding of the molecular and ecological factors driving the phage-bacteria interactions observed in this system. Extending such analyses to other ecologically and phylogenetically-distinct plant-associated bacteria, such as *Xanthomonas*, *Ralstonia*, or *Agrobacterium*, could reveal whether the patterns observed in *P. syringae*-phages infection network are generalisable or reflect unique evolutionary dynamics of this highly diverse species complex.

## Materials and Methods

### Phages panel

The 23 phages infecting strains of the *P. syringae* species complex used in this study were isolated from soil samples collected in apricot orchards affected by bacterial canker and fully described in a previous study (72). Briefly, this phage collection exhibits high genetic diversity, encompassing 13 genera and 17 species, representing the three most common morphological types (myovirus, podovirus, and siphovirus) and comprising 20 virulent and 3 temperate phages. The collection included two characterised DNA packaging mechanisms (direct terminal repeat (DTR) and headful) alongside two newly identified but unverified mechanisms, while the remaining packaging strategies remain unknown. The genomic GC content of the phages ranged from 46% to 58%. All phage stocks were standardised to a concentration of 10⁷ PFU/mL and stored in SM buffer (100 mM NaCl, 8 mM MgSO₄·7H₂O, 50 mM Tris-HCl, pH 7.4) at 4°C.

### Bacterial strains panel

The 44 *P. syringae* species complex strains were obtained from the Plant Pathology Unit collection at INRAE (Montfavet, France) and were not isolated as part of this project (S4 Table). The strains belonged to seven clades within four phylogroups (PG01, PG02, PG07, PG10) corresponding to the species levels, according to Berge *et al.* (60). The strains were sampled from three different environments: diseased apricot trees (exhibiting bacterial canker symptoms) (34), healthy plant tissues, and non-agricultural environments (S4 Table). Bacterial strains were revived from -80°C storage and cultured on King’s B agar (73). Cultures were maintained at 4°C for a maximum of one week.

### Sequencing and analysis of whole genomes of bacterial strains

The bacterial cultures were grown on King’s B (KB) agar medium for 48 hours at 25°C. A bacterial suspension was then prepared to achieve an approximate concentration of 2 × 10⁹ CFU in 4 mL of demineralised water. The suspension was subsequently centrifuged at 3,215 *g* for 10 minutes at 20°C to pellet the bacterial cells, after which the supernatant was carefully discarded. Genomic DNA was extracted using the Qiagen Genomic-tips kit (Qiagen, Valencia, CA, USA) following the manufacturer’s protocol.

Of the 44 *P. syringae* strains, 39 were sequenced using 20 µL of genomic DNA, with sequencing performed by Plasmidsaurus Inc (San Francisco, USA) using Oxford Nanopore Technology (ONT) and a custom analysis and annotation pipeline (https://plasmidsaurus.com/). The long-read sequencing library was prepared without amplification using v14 library preparation chemistry, and sequencing was conducted with R10.4.1 flow cells. The genomes of five additional strains were obtained from the RefSeq library at NCBI: CC1559 (GCF_000452685.1), USA0007 (GCF_000452545.1), 41A (GCF_000935775.1), MAFF302273^PT^ (also known as JD08) (GCF_000145925.1), and CC1583 (GCF_000452665.1).

The core genome analysis was performed using a private database within the EDGAR 3.5 platform (74). The core genome was defined as the set of orthologous genes present in all 332 genomes of the *P. syringae* species complex, including the 44 strains analysed in this study. A total of 1,346 core genes were identified per genome, corresponding to 446,872 aligned genes across all strains. Each set of orthologous genes was individually aligned at the protein level using MUSCLE (74). The resulting alignment were then concatenated into a single alignment, yielding approximatly 512,223 amino acid residues per genome and 170,058,036 in total across all genomes. Phylogenetic relationships were inferred using the FastTree algorithm within EDGAR, which constructs maximum-likelihood phylogenetic trees based on the core genome alignment (75). All phylogenetic trees of bacterial strains shown in the figures were rooted at the midpoint of the longest path and the nodes with a maximum likelihood support value below 0.90 were marked with red dots.

### Identification of defence systems and prophages in bacterial genomes

The defence systems of the 44 *P. syringae* strains were identified using *DefenseFinder* version 1.2.2 (76) based on assembled nucleotide sequences from the bacterial genomes. Each defence system type was categorised into one of three functional groups: nucleic acid degradation, abortive infection (Abi), or undetermined (ND).

The prophages within the 44 *P. syringae* strains were identified using *geNomad* version 1.5.0 based on assembled nucleotide sequences of bacterial genomes. As prophage prediction tools are still imperfect and may miss or misclassify certain elements, these results should be interpreted with caution. Among the 97 identified prophage sequences, only those classified as viral by *geNomad* and *CheckV* (78) were retained. Specifically, sequences labelled as “No Terminal Repeats” in the topology column and those with a “viral_genes” value of 0 were excluded. This filtering resulted in a final dataset of 88 verified viral prophages, accounting for 91% of the total identified prophages across the 44 strains. Additionally, a genetic analysis of the prophages was performed using *VIRIDIC* (79) to classify them into genera and species (S3 Table). *VipTree* was used to generate prophage clusters based on a proteomic tree analysis using distance-based methods (BIONJ from tBLASTx scores) (80). *geNomad* and *VipTree* were then employed to identify prophages belonging to the CL4 cluster and to construct a proteomic tree encompassing 68 CL4 prophages (24 reference strains + 44 analysed strains) (81).

To assess the congruence between the taxonomy of CL4 prophages and the phylogeny of their bacterial hosts, we performed a PACo (Procrustes Approach to Cophylogeny) analysis. Phylogenetic distance matrices were generated from the host and prophage trees using the *cophenetic* function (82). These matrices were then centred and corrected for negative eigenvalues using the “Cailliez” correction method. Congruence was tested using the *PACo* function from the *paco* R package with 1,000 permutations to assess statistical significance. The co-phylogeny visualisation was performed with *phytools* R package and *cophylo* function (83).

### Phage host range analysis

#### a. Experimental setup

To analyse the host range of phages infecting *P. syringae*, bacterial growth kinetics in the presence of phages were measured using 96-well plates (65). This liquid culture approach has been compared to other methods, including qPCR, spot assays, and plaque assays, demonstrating its relevance for estimating phage host range across diverse bacterial–phage combinations (65,84–86). Given the broad panel of phages and bacterial strains analysed, we selected this method as it provides an optimal balance between throughput, reproducibility, and comparability, while avoiding the more labour-intensive phage progeny quantification assays.

The dataset comprised 44 bacterial strains tested in biological triplicate without phage as a positive control, and the same 44 strains in biological triplicate exposed individually to each of the 23 phages (S6 Fig). Each phage was tested against each biological triplicate of bacterial strain, resulting in 3,036 growth kinetic measurements (23 phages × 44 bacterial strains × 3 replicates) (S6 Fig). Negative controls (culture medium) were used to check for contamination, with plates containing more than two contaminated wells out of four discarded (S6 Fig and S7A Fig). Two internal controls were included on each plate: the *P. syringae* strain P1.01.01B03 (PG01a, isolated from apricot buds in France in 2014) and phage Ghuch01, chosen because it completely inhibited the growth of strain P1.01.01B03 under the experimental conditions used in this study (34,72) (S6 Fig and S7B-C Fig). A plate was considered valid only when all three controls were visually confirmed: the internal bacterial control exhibited visible turbidity, while both the internal phage control and the media controls showed clear wells (S6 Fig).

Bacteria were incubated in 5 mL of liquid KB medium at 25°C and 200 rpm for 4 hours. Each replicate was standardised to an OD_600nm_ = 0.01 (∼10⁷ CFU/mL) in phosphate buffer and diluted in liquid KB medium to a final concentration of 10⁵ CFU/mL. Each 96-well spectrophotometric plate (NUNC, Thermo Scientific 267334) contained most phages confronted with a single bacterial strain, with positive control wells containing 180 µL of bacterial suspension and 20 µL of SM buffer. Experimental wells comprised 180 µL of bacterial suspension and 20 µL of phage suspension at 10⁷ PFU/mL, corresponding to a multiplicity of infection (MOI) of 10, as determined from preliminary optimization tests. Positive control measurements included between two and seven technical replicates per biological replicate (except for bacterial strains 41A and 93D, which had only one technical replicate per biological replicate). Measurements were taken using the SPECTROstar Omega spectrophotometer (BMG Labtech, France) every 5 minutes for 48 hours at 25°C, with intermittent orbital shaking at 200 rpm (200 seconds “ON” followed by 400 seconds “OFF”).

To assess the host specificity of the 23 phages, their infectivity was tested on four *Pseudomonas* spp. strains outside the *P. syringae* species complex using the double agar layer method on solid medium. The selected strains were *P. aeruginosa* CFBP 2466, *P. fluorescens* CFBP 2102, *P. graminis* 38B9, and *P. rhizosphaerae* 6B4. Strains were grown on King’s B (KB) agar for 48 h at 25°C (*P. fluorescens*, *P. graminis*) or 28°C (*P. aeruginosa*, *P. rhizosphaerae*). Biological triplicates were inoculated into 5 mL of KB broth and incubated overnight at 25°C with shaking at 200 rpm. Then, 500 µL of each culture was transferred to 5 mL fresh KB and incubated under the same conditions for 4 h. 300 µL of the culture in exponential phase (OD between 0.2 and 0.8 at 600 nm) was mixed with 6 mL of 0.7% KB semi-solid agar for the overlay. Each phage at an initial concentration of 10⁷ PFU/mL was serially diluted, and 10 µL of each dilution were spotted in duplicate onto the bacterial lawn. Plates were allowed to dry and incubated overnight at 25°C before evaluation of plaque lysis. The phage concentrations on the tested strains were divided by the concentration obtained on their respective reference strains to calculate the efficiency of plating (EOP), a standard metric.

#### b. Calculation of bacterial growth inhibition percentages and probability of significant inhibitory effects of phages on bacteria

Bacterial growth curve data from each plate were analysed using R software (version 4.4.0) with the *growthcurver* package (87). The area under the empirical growth curve (auc*e*) was the primary parameter analysed. For each biological replicate of the bacterial control condition (without phage), the mean and standard deviation of the auc*e* were calculated. The percentage of bacterial growth inhibition caused by each phage was then determined for each biological replicate using the following formula:

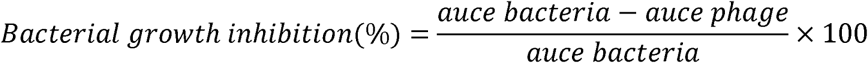

where auc*e* bacteria represent the mean auc*e* of each bacterial control biological replicates, and auc*e* phage corresponds to the auc*e* of the same bacterial strain when exposed to a given phage. This normalisation accounts for potential differences in bacterial growth rates, allowing for direct comparison of phage effects across bacterial strains (88). To evaluate potential bias in this calculation, particularly when bacteria exhibit different maximal densities (88), we assessed the coefficient of variation (CV) of the *k* parameter among bacteria control condition. The CV was low (9.22%), supporting the validity of this approach and indicating minimal bias associated with differences in growth maxima. Additionally, the CV of the auc*e* values under bacteria control condition was moderate (11.34%), further supporting the robustness of the percentage inhibition metric for cross-strain comparisons while limiting potential biases (88).

Phage impact on bacterial growth was assessed using a probability value derived from a normalised distribution, calibrated based on the mean (µ in the formula) and standard deviation (σ in the formula) of the auc*e* for each biological replicate of each bacterium exposed to phages. The probability ranges from 0 to 1 and is calculated using:

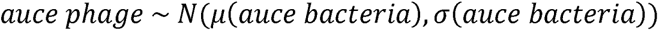

A probability value below 0.05 indicates a significant growth reduction in presence of phage, while a value exceeding 0.95 indicates a significant growth stimulation of bacteria in presence of phage. The distribution of inhibition percentages within the 0.05 to 0.95 probability range is approximately symmetric and unimodal (S8 Fig), resembling a Gaussian distribution. However, the Shapiro-Wilk test indicated a significant deviation from strict normality (p < 0.01), likely due to the large sample size. Despite this, the distribution shape supports the use of this probability window to robustly summarise phage effects on bacterial growth. Bacterial strains 41A and 93D had only one technical replicate per biological replicate, precluding probability calculation.

A phage was considered capable of inhibiting a bacterial strain if at least one of the three bacterial replicates showed a significant inhibition (probability < 0.05); in this case, the phage–bacterium combination was assigned a value of 1, in order to capture the maximal host range potential of each phage. If no replicates showed significant inhibition, the combination was assigned a value of 0. This threshold was verified to yield consistent results with a more stringent criterion (requiring at least two out of three replicates to be significant), in order to confirm the robustness of our analysis.

The relationship between the breadth of infection and the strength of inhibition was first examined using the full dataset. For the phage perspective, we plotted virulence (mean percentage of inhibition per phage) against infectivity (the number of bacterial strains each phage significantly infected) (S9A Fig). For the bacterial perspective, we plotted susceptibility (mean percentage of inhibition per bacterial strain) against sensitivity (the number of phages significantly infecting each strain) (S9B Fig). We then considered only combinations showing significant inhibition, for both phages (S9C Fig) and bacteria (S9D Fig), in order to reduce the dependency between variables. Spearman correlation tests were calculated exclusively on this subset of significant interactions, as this approach removes part of the inherent dependence generated by the shared underlying metric.

Generally, bacterial populations decreased in the presence of phages. In 17% of observed phage-bacteria combinations, a “stimulatory” effect was identified, with inhibition values reaching up to -40%. Given the lack of reproducibility of the stimulatory effect (S10 Fig), only positive inhibition values were retained for all analyses.

### Statistical analysis

#### a. Analysis of the structure of phage-bacteria interaction networks

The analysis of networks representing bacterial growth inhibition percentages of 44 bacterial strains combined with 23 phages was computed using the mean of the three biological replicates for each combination. The analyses were performed using the R software (version 4.4.0) with the *bipartite* (for nestedness and modularity analysis) and *igraph* (for modularity analysis) packages, following Moury *et al.* (30) and Torres-Barceló *et al.* (46).

The nestedness of the inhibition percentage network was evaluated using two algorithms: *wNODF* (a weighted nestedness metric based on overlap and decreasing fill) (89) and *WINE* (weighted-interaction nestedness estimator) (90). Modularity was assessed using six algorithms: *spinglass* (91–93), *edge betweenness* (91), *fast greedy* (94), *leading eigenvector* (95), *Beckett* (96), and *louvain* (97).

Prior to nestedness and modularity analyses, the continuous inhibitory percentage data of the matrix were transformed into discrete integer values spanning from 0 to 100.

In addition, we analysed a network of binary data (0 and 1 values) representing the statistical significance of bacterial growth inhibition by phages. For this network, three algorithms were used for nestedness: *WINE* as applied to quantitative data, *Binmatnest* (98), and *NODF2* (89) and three algorithms were used for modularity (*spinglass, leading eigenvector,* and *Beckett)*.

For each algorithm and each network, the calculated nestedness and modularity scores were compared with the scores obtained with seven (network of continuous inhibition data) or six (network of binary inhibition significance data) different null models (i.e. simulated networks for which no particular nestedness or modularity patterns are expected) in order to estimate the statistical significance of nestedness and modularity of the actual networks (99,30,100). Briefly, each null model differs by the constraints applied during randomization. The N model preserves the total sum and number of zeros but not row or column totals. C1 and R1 apply the same principle column- or row-wise, respectively. The B model maintains both row and column sums. The S model fully randomizes cell values, while C2 and R2 randomize values within columns or rows, respectively. The PD (Probabilistic Degree) model takes the original matrix and reshuffles positive infections while maintaining the same ‘degree distribution’, on average, of both phages and hosts while reshuffling. Together, these models span a gradient of constraint strength, from the least constrained (S, N) to the most constrained (B).

#### b. Statistical analyses to explain bacterial sensitivity and susceptibility to phages

All statistical analyses were conducted in R version 4.4.2. The phylogenetic signal of the number of phages infecting each bacterial strain was calculated using the *phylosig* function from the *phytools* package, based on a phylogenetic tree reconstructed from bacterial core-genome analysis (83).

The bacterial susceptibility profiles were tested with a PERmutational Multivariate ANalysis Of VAriance (PERMANOVA), a non-parametric method that assesses the influence of categorical variables on multivariate data by partitioning variation within a distance matrix. Unlike traditional ANOVA, it makes no assumptions about data distribution and relies on permutations to evaluate the statistical significance of group differences (101). Here, Bray– Curtis dissimilarity was used to construct a distance matrix based on the mean percentage of bacterial growth inhibition (from three replicates per combination). This matrix was then analysed using the *adonis2* function from the *vegan* R package, with 9,999 permutations to ensure robust p-value estimation (102). “*Phylogroup”, “ecological origin”,* and “*whether the strain was an isolation bacterial strain for the phages*” were included as explanatory variables. Multiple comparisons were conducted using the function *pairwise.adonis2* from the *pairwiseAdonis* package (103), which is specifically adapted for PERMANOVA post-hoc testing.

Additionally, the summed sensitivity, determined from the significant inhibition network (see above), was used as a response variable in a Generalised Linear Model (GLM) with a Poisson distribution with the function *glmmTMB* from the package *glmmTMB* (104). The model included “*phylogroup”*, “*ecological origin”*, and their interaction as fixed effects. As no significant interaction was detected, the simplified additive model (*phylogroup + ecological origin*) was retained. No overdispersion was observed in the data (deviance/degrees of freedom = 1.01). The effects of each explanatory variable were assessed using Type III ANOVA with the *Anova* function from the *car* package. Post-hoc pairwise comparisons were performed using Tukey’s test implemented via the *multcomp* and *multcompView* packages, with significance groups assigned accordingly.

#### c. Statistical analysis to explain infectivity and virulence of phages

The Phylogenetic Host Range Index (PHRI) (46), derived from the phylogenetic diversity index *Hp* (105), was calculated using the *hilldiv* package . For this analysis, the bacterial phylogenetic tree, constructed from core-genome analysis (see above), was transformed into an ultrametric tree, and the PHRI index was determined using the matrix of percentage inhibition.

The ranking of phage genera based on inhibition percentage (offset by +0.1 to avoid zeros) was assessed using a GLMM with a Gamma distribution with a log-link, suitable for strictly positive, continuous, and skewed data. The model included a random effect accounting for the “*phage”* tested, the “*bacterial”* strain tested, and “*replicates”* nested within each “*phage*:*bacterium”* combination. No overdispersion was detected (χ²/ddl = 0.67). Post-hoc pairwise comparisons between genera were performed using Tukey’s test.

The phage-host infection profiles were tested with a PERMANOVA test based on the distance matrix based on the mean percentage of bacterial growth inhibition compared with “*phage genus”* and “*whether the phage had been isolated using one of the bacterial strains included in the experimental panel*” explanatory variables (Bray-Curtis distance matrix; 9,999 permutations). Pairwise comparisons of phage infection patterns were performed using the *pairwise.adonis2* function (103).

#### d. Statistical analysis of defence systems and prophages

The phylogenetic signal of the total number of defence systems and the abundance of prophages was calculated using the *phylosig* function from the *phytools* package, based on a phylogenetic tree reconstructed from bacterial core-genome analysis (83).

The total number of bacterial defence systems per bacterial genome and the abundance of prophages were tested against the explanatory variables “*phylogroup”*, “*ecological origin”,* and “*growth rate”* in a GLM with a Poisson distribution with an additive model of the three explanatory variables. No overdispersion was observed in the data (deviance/degrees of freedom for defence systems = 0.61; for abundance of prophages = 0.47), and the effects of each explanatory variable were assessed using Type III ANOVA. Finally, the post-hoc pairwise comparisons were performed using Tukey’s test and significance groups were assigned accordingly.

The relationship between ANI of bacterial pairs and their percentage similarity in their defence systems (offset by +0.1 to avoid zeros) was assessed using a GLMM with a Gamma distribution and a log-link function. Random effects for each bacterium in the pair were included to account for the non-independence of observations. No overdispersion was detected (χ²/ddl = 0.24).

The defence system profiles, the proportions of the defence system categories, and prophages distribution profile across bacterial strains were analysed using a PERMANOVA test based on a Bray-Curtis dissimilarity matrix with 9,999 permutations. “*Phylogroup”*, “*ecological origin”*, and “*growth rate”* were included as explanatory variables. Pairwise comparisons of the defence system and prophage composition between groups were conducted using the *pairwise.adonis2* function (103).

Linear regressions were conducted to explain the number of phages infecting each bacterial strain, using the *total number of defence system subtypes per bacterium*, the *phylogroups*, and *ecological origins* as explanatory variables in a multiple linear regression model (adjusted R² = 0.21; p = 0.02 for the overall model). The same analysis was applied using the *number of prophages per bacterium* instead of defence systems, with *phylogroups* and *ecological origins* as predictors (adjusted R² = -0.06; p = 0.81 for the overall model). Before conducting these analyses, all assumptions of linear regression, including normality of residuals and homoscedasticity, were verified.

#### e. Statistical analysis of host range data with multimodel analysis

To account for the widest range of explanatory variables influencing the percentage of inhibition, we performed a multimodel analysis coupled with a generalised linear mixed model (GLMM). To enhance model efficiency, the number of explanatory variables was reduced through a multi-step process. Variables with more than 20% missing data were excluded. To ensure independence among explanatory variables before multimodel analysis, we assessed pairwise associations using Spearman’s correlation for quantitative variables, chi-squared tests for categorical variables, and Kruskal–Wallis tests for associations between categorical and quantitative variables. Variables showing significant correlations (p < 0.05) were considered dependent. In such cases, one of the correlated variables was retained, while the others were excluded. To further refine the selection, Principal Component Analyses (PCA) were conducted separately on the matrices representing the distributions of defence systems and prophages across bacterial genomes. The most informative variables contributing to the first principal components were retained for subsequent modelling.

The PCA on defence systems was performed without centring or scaling the data. Two dimensions accounted for 32.5% of the variance and led to the selection of the variables “*Septu”*, “*Gabija”*, and “*RloC”*, which were among the most frequent defence systems in the genomes. Similarly, the PCA on the distribution of prophage species in bacterial genomes, also conducted without centring or scaling (already binary), captured 14% of the variance in the first two dimensions and identified two perpendicular variable groups composed respectively of prophage species “*17”*, “*54”*, “*26”,* and “*29”* on one side, and species “*7”*, “*47”*, “*57”*, “*22”*, and “*31”* on the other. One representative species from each group was selected for the multimodel analysis (species “*17”* and “*22”*).

To identify factors explaining variations in bacterial growth inhibition percentage, we fitted a GLMM. The response variable (offset by +0.1 to avoid zeros) was modelled using a Gamma distribution with a log-link function. Random effects included “*phage”*, “*bacterium”*, and their “*imbricated replicates”*.

To compare two approaches for modelling the effects of bacterial defence systems and prophages on the percentage of inhibition, two separate GLMMs were constructed: one using total counts of defence systems and prophages, and another incorporating PCA-derived principal components of the distribution matrices (of prophages and defence systems). No overdispersion was detected in either model (total counts: χ²/df = 0.65; PCA-derived: χ²/df = 0.66), supporting the appropriateness of the Gamma distribution for modelling the response variable.

To identify the most informative predictors explaining variation in phage replication efficiency, we used a multimodel analysis. Model selection was performed using the *dredge* function from the *MuMIn* package, ranking models based on corrected Akaike information criterion (AIC). All possible sub-models were generated and evaluated. Models whose cumulative AIC weights accounted for 95% of the total were retained for model averaging using the *model.avg* function. The relative importance of each explanatory variable was then estimated using the *sw* function, which computes the sum of the AIC weights across all models in which the variable appears.

## Acknowledgments

We thank Elisabeth Villard, Cécile Monteil, and Charlotte Chandeysson for their assistance with experimental procedures. We are also grateful to Dominique Holtappels for valuable discussions, and to Hannah Toth for her assistance in depositing bacterial genomes into NCBI.

## Data Availability Statement

The phage–bacterium infection network, based on average values, has been deposited in the Viral Host Range Database hosted by Institut Pasteur (https://viralhostrangedb.pasteur.cloud/data-source/287/). The raw optical density (OD) kinetics data and the processed datasets used for analysis (e.g., AUC, inhibition values, phage effect probability) have been deposited in Zenodo (DOI: 10.5281/zenodo.17735985). The 39 bacterial genomes sequenced in this study have been deposited under the BioProject accession number PRJNA1292126.

